# Genomes of gut bacteria from *Nasonia* wasps shed light on phylosymbiosis and microbe-assisted hybrid breakdown

**DOI:** 10.1101/2021.02.13.431100

**Authors:** Karissa L. Cross, Brittany A. Leigh, E. Anne Hatmaker, Aram Mikaelyan, Asia K. Miller, Seth R. Bordenstein

## Abstract

Phylosymbiosis is a cross-system trend whereby microbial community relationships recapitulate the host phylogeny. In *Nasonia* parasitoid wasps, phylosymbiosis occurs throughout development, is distinguishable between sexes, and benefits host development and survival. Moreover, the microbiome shifts in hybrids as a rare *Proteus* bacteria in the microbiome becomes dominant. The larval hybrids then catastrophically succumb to bacterial-assisted lethality and reproductive isolation between the species. Two important questions for understanding phylosymbiosis and bacterial-assisted lethality in hybrids are: (i) Do the *Nasonia* bacterial genomes differ from other animal isolates and (ii) Are the hybrid bacterial genomes the same as those in the parental species? Here we report the cultivation, whole genome sequencing, and comparative analyses of the most abundant gut bacteria in *Nasonia* larvae, *Providencia rettgeri* and *Proteus mirabilis*. Characterization of new isolates shows *Proteus mirabilis* forms a more robust biofilm than *Providencia rettgeri* and when grown in co-culture, *Proteus mirabilis* significantly outcompetes *Providencia rettgeri. Providencia rettgeri* genomes from *Nasonia* are similar to each other and more divergent to pathogenic, human-associates strains. *Proteus mirabilis* from *N. vitripennis, N. giraulti*, and their hybrid offspring are nearly identical and relatively distinct from human isolates. These results indicate that members of the larval gut microbiome within *Nasonia* are most similar to each other, and the strain of the dominant *Proteus mirabilis* in hybrids is resident in parental species. Holobiont interactions between shared, resident members of the wasp microbiome and the host underpin phylosymbiosis and hybrid breakdown.

**IMPORTANCE:** Animal and plant hosts often establish intimate relationships with their microbiomes. In varied environments, closely-related host species share more similar microbiomes, a pattern termed phylosymbiosis. When phylosymbiosis is functionally significant and beneficial, microbial transplants between host species or host hybridization can have detrimental consequences on host biology. In the *Nasonia* parasitoid wasp genus that contains a phylosymbiotic gut community, both effects occur and provide evidence for selective pressures on the holobiont. Here, we show that bacterial genomes in *Nasonia* differ from other environments and harbor genes with unique functions that may regulate phylosymbiotic relationships. Furthermore, the bacteria in hybrids are identical to parental species, thus supporting a hologenomic tenet that the same members of the microbiome and the host genome impact phylosymbiosis, hybrid breakdown, and speciation.

## INTRODUCTION

Microbiomes can play pivotal roles in host health and disease [1–6] and frequently establish distinguishable associations with host lineages [7–14]. Often the evolutionary relationships between closely-related host species mirror the ecological relationships of their microbial communities in a pattern termed phylosymbiosis [15]. Across animal and plant clades, closely-related host species harbor more similar microbial communities than divergent host species [9, 15–20]. Breakdown of phylosymbiotic relationships can also occur when the host genome and microbiome, or hologenome, is altered, such as in hybrid hosts [21–23] or parental hosts receiving a microbiome transplantation from another species [9, 24]. It can lead to detrimental effects including hybrid lethality and intestinal pathology [21, 22, 24, 25]. Resultantly, it is pertinent to investigate how the bacterial identities present within parental species are or are not altered as a result of hybridization.

A well-developed animal system for studying phylosymbiosis and hologenomic speciation is the *Nasonia* parasitoid wasp genus. It is comprised of four species whose ancestor arose approximately one million years ago (MYA) [26]. These species are interfertile in the absence of *Wolbachia* endosymbionts and exhibit strong trends in phylosymbiosis even under identical rearing conditions [7, 9, 27]. Crosses between *Nasonia* species long cured of their intracellular *Wolbachia* endosymbionts readily produce fit F_1_ hybrid females, but most of the F_2_ hybrid males die during larval development due to microbe-assisted hybrid lethality that occurs in association with changes in the microbiome, hypermelanization, and transcriptome-wide upregulation of immune gene expression [21, 28]. Moreover, transplants of microbiomes into larvae of each *Nasonia* species elicits reductions in host developmental rates and survival, supporting the premise that selective pressures drive phylosymbiosis in the system [24].

Early in development, *Nasonia* larvae possess a simple microbiome that changes composition throughout development [27]. Larvae of the two most divergent species, *N. vitripennis* and *N. giraulti*, possess gut microbiomes dominated by *Providencia* spp. bacteria (81-96% of sequencing reads), whereas their F_2_ larval hybrid offspring are dominated by *Proteus* spp. bacteria (86% of sequencing reads) [21]. Furthermore, these F_2_ interspecific hybrids exhibit ∼90% hybrid lethality between the third and fourth larval instars [21], and germ-free rearing can remarkably rescue the hybrid lethality [21, 29]. The rescue of hybrid lethality via germ-free rearing and recapitulation of death upon inoculation strongly supports the dependency of hybrid lethality on gut bacteria [30]. Interspecific microbiome transplants between *Nasonia* species with heat-killed bacterial communities also results in slowed larval growth and decreased pupation and adult survival, demonstrating how host responses play an integral role in interacting with their microbiomes [24]. The costs that result from gut microbiome transplants occur in an evolutionary-informed manner, which further suggests that selective pressures can underpin phylosymbiosis and holobiont composition [24].

The genomes of both *N. vitripennis* and *N. giraulti* are published [31, 32], and thus the imperative turns to the bacterial genomes within these two species and their hybrid offspring to understand the nature of the catastrophic events that result in gut bacteria-assisted hybrid lethality. In particular, are the bacteria in *Nasonia* guts distinguishable from other environmental isolates and thus specific to the species complex? And upon hybridization and phylosymbiotic breakdown, are the gut bacteria in parental hosts identical or different to the bacteria in hybrid hosts? A key aspect in this system is whether the same bacteria present in parental species contribute to reproductive isolation in hybrids?

In *Nasonia*, an alteration in the abundances of *Proteus* and *Providencia* bacteria in hybrid offspring associates with lethality [21]. *Proteus* and *Providencia* are well-characterized bacterial genera due to their role as opportunistic pathogens in both humans and insects [33–36]. *Proteus* spp. are present in low abundance in humans, but their overgrowth is often associated with urinary tract infections [35, 37]. In *Drosophila melanogaster*, different strains of *Providencia* display varying degrees of virulence towards the host whereby the host develops an immune response to potentially combat bacterial infection [38–40]. Additionally, in *Caenorhabditis elegans*, commensal *Providencia* bacteria in the gut produce a neurotransmitter that promotes fitness of both the host and bacteria [41]. The evolution of varying host responses to different bacteria demonstrates how intimate interactions between these bacteria and their hosts may mediate pathogenic versus symbiotic relationships.

Since *Proteus* and *Providencia* bacteria are native members of the *Nasonia* gut microbiome across multiple lines [27, 42], there are several questions as to whether or not changes in the strains of gut bacteria mediate hybrid lethality and whether the strains in *Nasonia* are distinct from other animals. As *Proteus* and *Providencia* bacteria are readily culturable in the laboratory, we isolated and sequenced the genomes of representative species from *Nasonia* and their hybrid larvae to investigate their genomic diversity of (i) *Proteus* and *Providencia* isolates between *Nasonia* and publicly available genomes from insects and humans, (ii) *Proteus* and *Providencia* isolates between parental *N. vitripennis* and *N. giraulti* species, and (iii) *Proteus* isolates between parental and F_2_ hybrid offspring to determine if they are the same. Annotation and evolutionary-guided comparisons of the gene content may inform the nature of what drives phylosymbiosis in *Nasonia* from the microbial side and what is the nature of bacterial-dependent hybrid lethality.

## RESULTS AND DISCUSSION

### Biofilm formation and genomics of *Providencia* and *Proteus* bacterial isolates from *Nasonia* spp. and their hybrids

We previously found that the dominant bacterial genus present in the microbiome of the larvae of *N. giraulti* and *N. vitripennis* is the genus *Providencia*, whereas in F_2_ recombinant male hybrids of the same developmental stage, the bacterial genus *Proteus* becomes the dominant taxa associated with severe hybrid mortality [21, 27]. Both of these bacteria are easily cultivable, well-studied in arthropods, and opportunistic human pathogens [6, 33–35, 43, 44]. Therefore, we sought to isolate and sequence the genomes of representative *Proteus* and *Providencia* species from parental *N. vitripennis* or *N. giraulti* and F_2_ hybrids derived from the cross of *N. vitripennis* males x *N. giraulti* females. We concurrently set up intra- and interspecific crosses and collected F_2_ third instar larvae (L3) from virgin F_1_ females. F_2_ male larvae were surface sterilized with 70% ethanol and sterile water and then homogenized. Homogenate was serially diluted on Tryptic Soy Agar (TSA) plates, and distinct bacterial colonies were randomly chosen, sub-cultured, and sent for whole genome sequencing **(Figure 1a)**.

**Figure 1.**
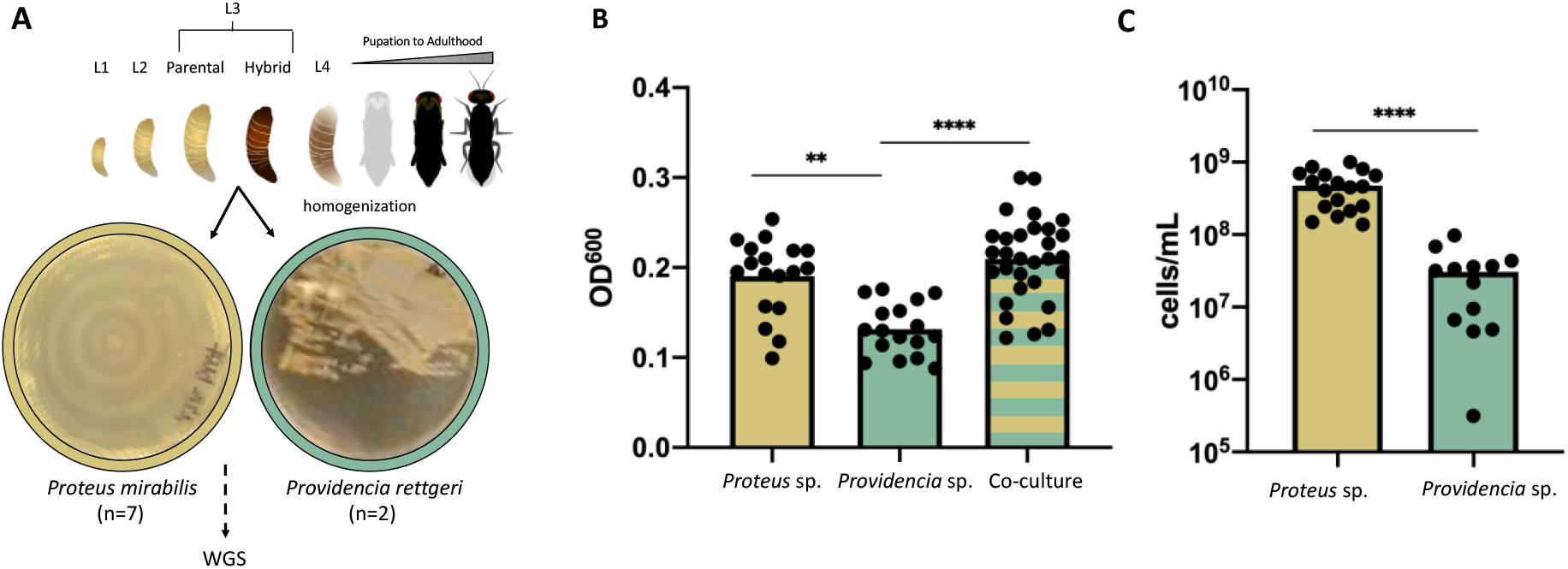
Characteristics of *Proteus mirabilis* and *Providencia rettgeri* isolates. **A)** Workflow for sample collection and examples of unique colony morphologies of *Providencia* (n=2) and *Proteus* (n=7) isolates. *Nasonia* development from larvae to adulthood is depicted on top demonstrating the L3 larval stage in which samples were collected from parental and hybrid lines. Following isolation, samples were prepared for whole genome sequencing (WGS). **B)** Biofilm formation by representative *Proteus* and *Providencia* isolates show significantly different abilities to form biofilms on solid surfaces. Bars denote significantly different values based on Kruskal-Wallis test with Dunn’s multiple comparison with sample size of n=17-30 of two biological replicates (*\*\* p*=0.0018; *\*\*\*\** p=<0.0001). **C)** Composition of the biofilm co-culture shows *Proteus mirabilis* significantly outcompetes *Providencia rettgeri* when grown together. *Providencia rettgeri* makes up about 6.4% of the co-culture biofilm composition. Bars denote significantly different values calculated by a Mann-Whitney U test with sample size of n = 13-18 (*\*\*\*\** p=<0.0001).

Bacteria that successfully colonize insect guts often adhere to surfaces and rapidly form biofilms [45]. To assess the ability for *Proteus* and *Providencia* to adhere and form biofilms on solid surfaces, we plated representatives from each genus in single culture and co-culture, and we measured their resulting biofilms. Over 24 hours, *Proteus mirabilis* formed a more robust biofilm than *Providencia rettgeri*, and when both isolates were plated together in a 1:1 ratio, the biofilm was significantly more abundant than *Providencia rettgeri* alone **(Figure 1b)**. To determine whether the slight, insignificant biofilm increase in the co-culture occurred in an additive manner with both species contributing equally to the biofilm or if the increase was principally due to one of the two species, we developed a qPCR assay to quantify their abundance. *Proteus mirabilis* significantly outcompeted *Providencia rettgeri* when grown together (*p*<0.0001) with *Providencia rettgeri* only making up about 6.4% of the biofilm composition **(Figure 1c)**. This suggests that *Nasonia* host genetic factors may keep the *Proteus* bacterial abundances under control *in vivo* in parental wasp species; that regulation is then compromised in hybrids, where *Proteus* dominates the microbiome or when the two bacteria are grown in coculture outside of the host. *Proteus mirabilis* is a well-characterized human pathogen [46] that has a distinctive swarming behavior which serves to reduce competition for nutrients between bacterial strains [47]. When *Nasonia* hybridization produces a F_2_ genotypic recombinant that shuffles the genes of the two host species, the swarming behavior of *Proteus mirabilis* coupled with a presumptive breakdown in the ability of the host to regulate bacterial abundances may create an environment that permits bacterial overgrowth and ensuing host lethality.

Following isolation of colonies and DNA extraction, whole genome sequencing was performed using an Illumina MiSeq. Data on genome statistics including number of reads generated, sequencing coverage, and genome size and completion are provided in Tables 1 and 2. For strain nomenclature, Ngir or G, Nvit or V, and Hybrid or H denote *N. giraulti, N. vitripennis*, and hybrid, respectively; “L3”, if noted, represents the third larval instar stage used for bacterial isolation and cultivation; numbers at the end of the name are unique designations assigned upon isolation (#1-3). Shorthand notation of all genomes presented throughout this study are provided in Supplemental Table 1. Isolate identity was confirmed after sequencing by comparison of the 16S rRNA sequences extracted from each genome to the RefSeq database.

**Table 1.** *Providencia rettgeri* sequencing and genome statistics.

| Strain Reference | Wasp.NgirL3.3 | Wasp.NvitL3.3 | Wasp.Nvit03 | Fly.Dmel1 | Human.RB151 |
| --- | --- | --- | --- | --- | --- |
| NCBI Strain Name | NgirL3-3G | NvitL3-3V | NVIT03<sup>†</sup> | Dmel1<sup>†</sup> | RB151* |
| Host Source | *N. giraulti*<br>L3 Larvae | *N. vitripennis*<br>L3 Larvae | *N. vitripennis*<br>gut | *D. melanogaster*<br>hemolymph | Human<br>(urine UTI) |
| Reads | 3,774,124 | 1,321,295 | 191,241 | 680,000 | 167,518 |
| Contigs | 16 | 20 | 73 | 9 | 1 |
| Average Coverage | 163 | 189 | 164 | 30 | 581 |
| Genome Size | 4,254,678 bp | 4,314,756 bp | 4,397,852 bp | 4,251,508 bp | 4,780,676 bp |
| GC content | 40.2 | 40.2 | 40.3 | 40.2 | 40.5 |
| CDS (coding) | 3,898 | 3,897 | 4,076 | 3,848 | 4,484 |
| Plasmids | 1 | 1 | 0 | 1 | 1 |
| Prophage regions | 3 | 3 | 6 | 4 | 2 |
| % Completed | 100 | 99.91 | 100 | 98.61 | 99.46 |
| % Contaminated | 1.62 | 1.89 | 1.08 | 1.08 | 1.08 |
| Reference | This study | This study | Wang GH,<br>et al. 2019 | Galac MR,<br>et al. 2012 | Marquez-Ortiz RA,<br>et al. 2017 |

<sup>†</sup> Previously published insect isolates (*Accession: NVIT03 QUAF01000007; Dmel1 CM001774*)
\* Human isolate reference strain (*Accession: RB151 PRJNA342046*)

**Table 2.** *Proteus mirabilis* sequencing and genome statistics.

| Strain Reference | Wasp.NgirL3.1 | Wasp.NgirL3.2 | Wasp.HybL3.1 | Wasp.HybL3.2 | Wasp.HybL3.3 | Wasp.NvitL3.1 | Wasp.NvitL3.2 | Human.HI4320* |
| --- | --- | --- | --- | --- | --- | --- | --- | --- |
| NCBI Strain Name | NgirL3-1G | NgirL3-2G | F<sub>2</sub>Hybrid-1H | F<sub>2</sub>Hybrid-2H | F<sub>2</sub>Hybrid-3H | NvitL3-1V | NvitL3-2V | HI4320* |
| Host Source | *N. giraulti*<br>L3 Larvae | *N. giraulti*<br>L3 Larvae | *Nasonia*<br>L3 Hybrid | *Nasonia*<br>L3 Hybrid | *Nasonia*<br>L3 Hybrid | *N. vitripennis*<br>L3 Larvae | *N. vitripennis*<br>L3 Larvae | Human<br>(urine) |
| Reads | 4,180,770 | 4,215,774 | 3,928,914 | 3,823,446 | 3,926,886 | 4,193,134 | 4,059,594 | 60,288 |
| Contigs | 57 | 61 | 46 | 62 | 63 | 42 | 62 | 1 |
| Average coverage | 399 | 348 | 443 | 323 | 331 | 471 | 377 | 8 |
| Genome Size | 4,024,338 bp | 4,023,051 bp | 4,076,851 bp | 4,023,528 bp | 4,024,027 bp | 4,070,909 bp | 4,026,776 bp | 4,063,606 bp |
| GC content | 38.6 | 38.6 | 38.6 | 38.6 | 38.6 | 38.6 | 38.6 | 38.8 |
| CDS (coding) | 3,639 | 3,632 | 3,688 | 3,621 | 3,628 | 3,683 | 3,628 | 3,611 |
| Plasmids | 0 | 0 | 0 | 0 | 0 | 0 | 0 | 1 |
| Prophage regions | 19 | 17 | 19 | 19 | 17 | 18 | 17 | 4 |
| % Completed | 100 | 100 | 100 | 100 | 100 | 100 | 100 | 100 |
| % Contaminated | 0 | 0 | 0 | 0 | 0 | 0 | 0 | 0 |
| Reference | This study | This study | This study | This study | This study | This study | This study | Pearson MM, et al. 2008 |

\* Human isolate reference strain (*Accession: HI4320 PRJNA342046*)

We sequenced *Providencia rettgeri* isolates from *N. giraulti* and *N. vitripennis* parental lines. The genome size for *Providencia* strain Wasp.NgirL3.3 is 4,254,678 bp with 163-fold coverage; and Wasp.NvitL3.3 is 4,324,254 bp with 189-fold coverage. Both strains have a G+C content of 40.2%. A single plasmid of 75,229 bp was identified in *Providencia* strain Wasp.NgirL3.3 with 324-fold coverage, and a single plasmid of 75,094 bp was identified in Wasp.NvitL3.3 with a 116-fold coverage. Both plasmids have 100% nucleotide identity between them.

The average genome size for the seven *Proteus mirabilis* isolates is 4,046,931 bp ranging between 3,823,446 to 4,215,774 bp with an average coverage of 385-fold. The G+C content of all *Proteus mirabilis* isolates is 38.6%. No circular elements were identified in any isolate. Using CheckM [48] for genome completion and contamination estimates, the *Providencia rettgeri* and *Proteus mirabilis* genomes are predicted to be 99.91-100% complete with 0.00-1.89% contamination (Tables 1 and 2).

### Phylogenetic placement of *Nasonia* bacterial isolates with other host-associated lineages

To characterize and compare the genomic diversity of the sequenced *Providencia rettgeri* and *Proteus mirabilis* isolates, we surveyed the National Center for Biotechnology Information (NCBI) databases for representative genomes. *Proteus mirabilis* is a commensal, gastrointestinal bacteria in host-associated environments [46, 49], including in humans [50], tree shrews [51, 52], and insects [21, 27, 53]. It is well characterized due to its role as an opportunistic pathogen in different clinical settings [33–36], especially urinary tract infections involving catheters [37] as well as bacteremia and wound infections [46]. However, less is known regarding *Providencia rettgeri*, despite also serving as an opportunistic pathogen in both humans and insects [5, 6, 43, 44, 54]. We characterized our *Nasonia* isolates based on single copy gene phylogenies of the housekeeping genes *DNA gyrase B* (*gyrB*, topoisomerase) and *RNA Polymerase beta subunit* (*rpoB*, RNA synthesis), a 70-71 multi-protein concatenated phylogeny **(Figure 2)**, and whole genome average nucleotide identity (ANI) for all-against-all pairwise comparisons with publicly available whole genome sequences **(Supplemental Figure 1)**. We report two key results.

**Figure 2.**
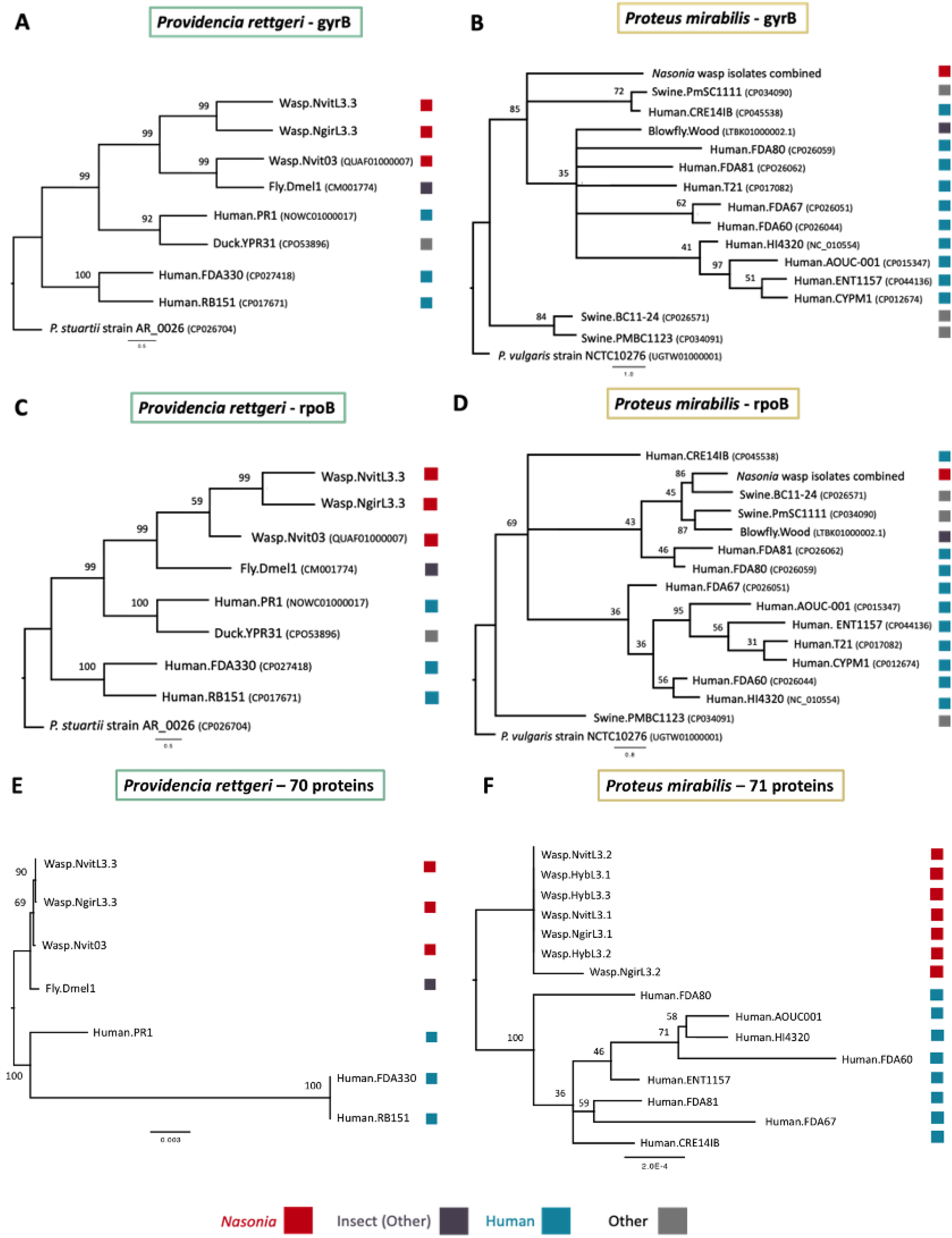
Phylogenetic placement of *Nasonia* isolates relative to publicly available genomes. **A-B)** Phylogeny of *Providencia rettgeri* and *Proteus mirabilis* strains based on 2,415 nucleotides of the gyrase B subunit reconstructed using RAxML. **C-D)** Phylogeny of *Providencia rettgeri* and *Proteus mirabilis* strains based on 4,029 nucleotides of the RNA Polymerase beta subunit reconstructed using RAxML. **E-F)** Phylogeny of *Providencia rettgeri* and *Proteus mirabilis* strains used in this study for comparative genomics analyses (Supplemental Table 1) based on concatenated amino acid alignments of 70-71 core bacterial proteins reconstructed using PhyML.

First, the *Providencia rettgeri* insect isolates are distinct from human-associated isolates. In the maximum likelihood (ML) phylogeny for both *rpoB* and *gyrB*, the *Nasonia* isolates from this study form a well-supported clade with previously-published, insect-associated *Providencia rettgeri* strains Wasp.Nvit03 (QUAF01000007) from *Nasonia vitripennis* [42] and Fly.Dmel1 (NZ_CM001774) from *Drosophila melanogaster* [54] **(Figure 2a,c)**. To explore these relationships further, we also built a concatenated protein tree using 70 core bacterial proteins to garner a higher resolution view of the relatedness between *Providencia rettgeri* strains used in this study **(Supplemental Table 1)**. A split in the phylogeny of these strains was apparent based on host origin, whereby *Nasonia* isolates grouped together with *D. melanogaster* and apart from human isolates, consistent with the ML phylogenies **(Figure 2e)**. Lastly, based on whole genome pairwise average nucleotide identity (ANI) [55], we classified Fly.Dmel1, Wasp.Nvit03, Wasp.NvitL3.3, and Wasp.NgirL3.3 as the same bacterial strain based on their >99% ANI. The closest human reference isolate to the insect *Providencia rettgeri* isolates was strain Human.PR1 with an ANI of ∼92% followed by strains Human.RB151 and Human.FDA330 at an ANI of ∼84% **(Supplemental Figure 1)**. ANI values >=95% are considered the same species [55, 56]; therefore, the human isolates of *Providencia rettgeri* may represent different species.

The *Proteus mirabilis* isolates sequenced from *Nasonia* have identical *gyrB* and *rpoB* sequences at the nucleotide level, and a representative sequence was used to build the ML phylogenies **(Figure 2b,d)**. The *Proteus mirabilis* ML trees **(Figure 2b,d)** provide moderate support for the placement of the *Nasonia* isolates, with most publicly available human-associated *Proteus mirabilis* genomes forming a separate clade with low support. The low phylogenetic support is due to the high similarity of these genes between close relatives. Therefore, we built a concatenated amino acid protein tree using 71 core bacterial genes to provide more resolution to the diversity within the *Proteus mirabilis* isolates **(Figure 2f, Supplemental Table 1)**. The phylogenomic tree showed that *Nasonia*-associated *Proteus mirabilis* are nearly 100% identical and form a distinct grouping from human-associated strains. Lastly, based on ANI, all *Proteus mirabilis* isolates from *Nasonia* were identified as the same strain with >99% ANI. When comparing human-associated whole genome sequences of *Proteus mirabilis* with our *Nasonia* isolates, all isolates are classified as the same subspecies with an ANI >98.5% (**Figure 2b,d**; **Supplemental Figure 1**). Collectively, there is little to no genetic diversity present amongst all the *Proteus mirabilis* bacterial isolates from *Nasonia* species and their hybrids.

### Comparative genomics within and between bacteria from *Nasonia* and other animals

#### *Providencia rettgeri* pangenomics

For comparative genomic analyses, we utilized five publicly available high-quality whole genome sequences of *Providencia rettgeri* to investigate the unique genomic changes that arose in *Nasonia-*associated and other animal-associated *Providencia rettgeri* isolates. The *Providencia rettgeri* genomes are provided in Supplemental Table 1. The public genomes are from *Nasonia vitripennis* (Wasp.Nvit03) [42], *Drosophila melanogaster* (Fly.Dmel1) [54], and three human samples (reference strain Human.RB151 [44], Human.FDA330 [57], and Human.PR1). Combined with our two *Nasonia*-associated isolates of *Providencia rettgeri*, Wasp.NvitL3.3 and Wasp.NgirL3.3, pangenomic analyses in anvi’o [58, 59] produced a total of 6,789 gene clusters (COGs). We grouped these gene clusters into seven bins based on their presence/absence across the genomes: (i) *Providencia* spp. Core (2,949 gene clusters), (ii) Human Specific (58 gene clusters), (iii) *Nasonia* Specific (55 gene clusters), (iv) Insect Specific (142 gene clusters), (v) *Drosophila* Specific (197 gene clusters), (vi) *Nasonia* – Nashville (168 gene clusters), and (vii) *Nasonia* – Cambridge (269 gene clusters) **(Figure 3)**. The city nomenclature between isolates Wasp.NvitL3.3, Wasp.NgirL3.3, and Wasp.Nvit03 is used to denote potential genomic differences between laboratories and geographies as these two lines were derived from the same *N. vitripennis* line less than six years apart and reared apart since then. More genomes from these lines will be required to further evaluate the evolution of genomic differences. The core genome represents gene clusters present in all genomes analyzed (human-associated and insect-associated). Most gene clusters (43.4%) fall within the *Providencia* spp. Core bin, as expected, where the majority of COG functions relate to [C] Energy production and conversion, [E] Amino acid transport and metabolism, and [J] Translation, ribosomal structure and biogenesis **(Supplemental Table 2, 3)**.

**Figure 3.**
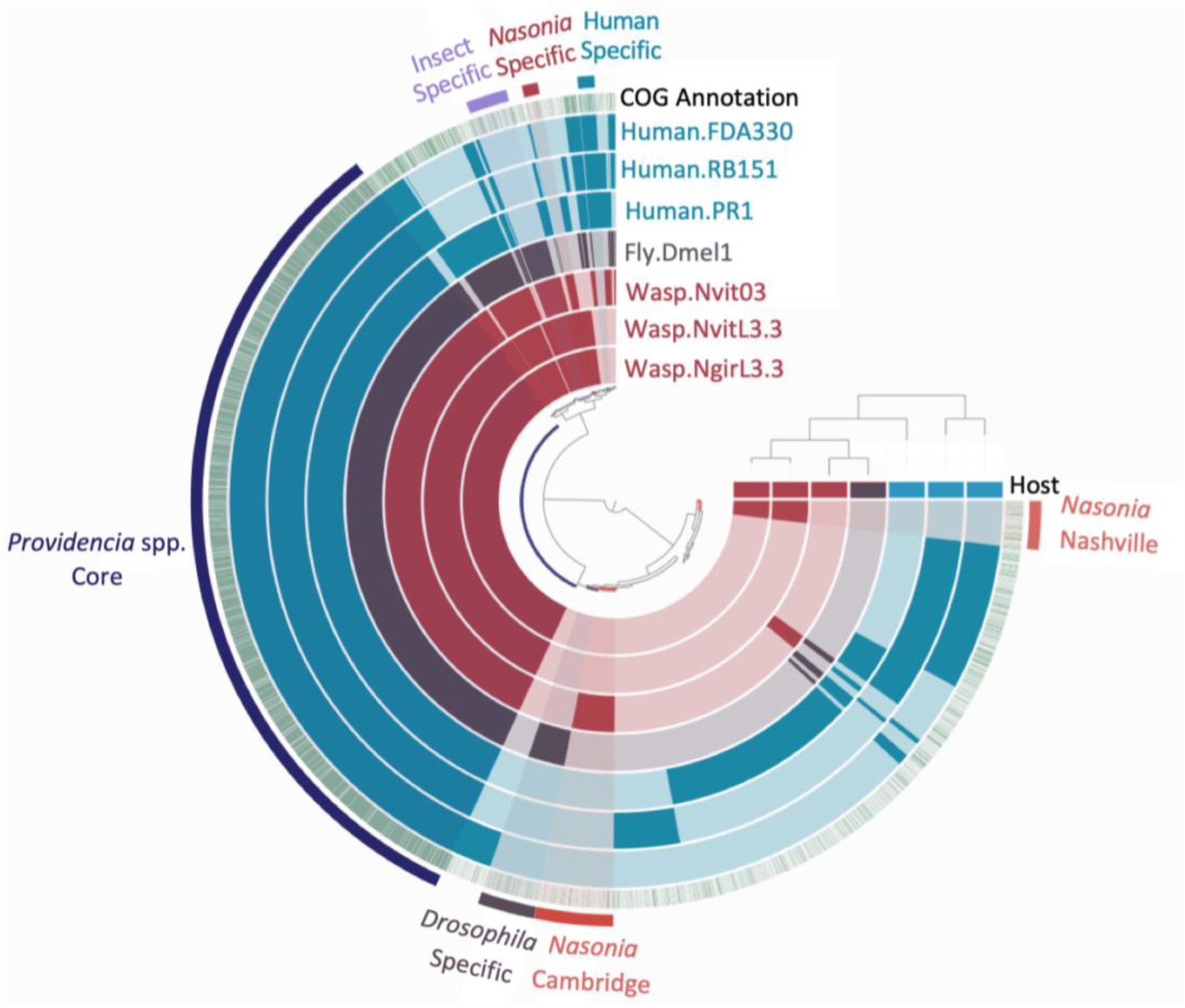
Pangenome of *Providencia rettgeri* human and insect isolates. The seven inner layers correspond to the seven genomes, including the sequenced *Nasonia* isolates in maroon, previously published *Drosophila* isolate in purple, and human associated genomes in blue. The bars in the inner circles show the presence of gene-clusters (GCs) in a given genome with the outer green circle depicting known COGs (green) vs unknown (white) assignments. The outermost layer of color-coded lines and text highlight groups of GCs that correspond to the genome core or to group-specific GCs. Genomes are ordered based on their gene-cluster distribution across genomes, which is shown at the top right corner (tree). Central dendrograms depict protein cluster hierarchy when displayed as protein cluster frequency. The top horizontal layer underneath the tree represent host source (*Nasonia -*maroon, *Drosophila -* purple, or Human - blue).

As aforementioned, we determined the phylogenetic relationships among these *Providencia rettgeri* whole genome sequences and noted a split in the phylogeny of these strains based on host origin, which is consistent with their ANI divergence **(Figure 2, Supplemental Figures 1)**. As adaptations to specific host environments may drive strain divergence in *Providencia rettgeri*, we sought to investigate host-associated genomic variation in *Providencia rettgeri* between (i) humans and insects, (ii) *Nasonia* and human/*Drosophila*, and (iii) various *Nasonia* isolates to further explore how bacterial genomic variation partitions across host variation.

#### Functional differences between Providencia rettgeri from insects and humans

The human and insect bins encompass accessory proteins, also referred to as gene clusters (GCs), present in either all human- or insect-associated genomes (*Nasonia* and *Drosophila* combined) **(Figure 3)**. We divide the discussion of these unique bins to those with shared functions and then those without shared functions. Distinct gene clusters between human and insect isolates based on the Markov Cluster Algorithm (MCL) [60, 61] that share homologous functional assignments (defined as distinct proteins having the same COG assignment) span hemolysins, large exoproteins involved in heme utilization or adhesin, and adhesin and type 1 pilus proteins **(Supplemental Table 2)** These functions are typically involved in host cell lysis [62], biofilm formation [63], and attachment to mucosal surfaces [63–65]. Notably, COG3539 for pilin (type 1 fimbria component protein) is the most abundant functional annotation in all *Providencia rettgeri* genomes, averaging around 58 occurrences in each genome. Different proteins encoding similar functions between human and insect-associated bacterial genomes could be due to slightly different adaptations of these genes as the bacteria evolved within their specified niche.

Next, we investigated what functions are present in the human-associated *Providencia rettgeri* genomes and missing or reduced in insect-associated isolates. In the human-associated *Providencia rettgeri* genomes, there are twice as many proteins relating to COG functions for [U] intracellular trafficking, secretion, and vesicular transport (7.05% vs. 2.35% of total COG functions) **(Supplemental Table 2, 3)**. This difference seems to be primarily driven by the increased occurrence of proteins relating to some of the same functions above, namely hemolysins and large exoproteins involved in heme utilization or adhesin. Large exoproteins involved in heme utilization or adhesin (COG3210) are secreted to outer bacterial membranes and serve as attachment factors to host cells [65]. For example, *Providencia stuartii* expressing higher levels of MR/K hemagglutinin adhered better to catheters and led to persistence in catheter-associated bacteriuria [66]. Similarly, the filamentous hemagglutinin (FHA) protein secreted by *Bordetella pertussis* is used for attachment to the host epithelium early in its pathogenesis [67]. Collectively, the increased presence of these genes in human-associated *Providencia rettgeri* suggests an adaptive lifestyle towards human colonization and virulence. Related, *Drosophila* species have developed unique antimicrobial peptides that suppress and resist *Providencia rettgeri* by reducing bacterial burden [39, 40].

Furthermore, CRISPR-Cas system-associated endonucleases Cas1 and Cas3 were strictly in human-associated *Providencia rettgeri* strains Human.FDA330 and Human.RB151 and absent in all other strains. The adaptive CRISPR-Cas systems are multigenic and include short repeated palindromic regions alongside spacers of phage DNA that serve as recognition factors triggering an “immune” response when phages encoding the spacer attempt to infect the microbe [68]. CRISPR-Cas systems are crucial for subversion of phage, and as such, human-associated isolates with CRIPSR-Cas have fewer integrated prophages (five total; n=2) than the *Nasonia*-associated ones (12 total; n=3) **(Table 1)**. The human-associated isolate Human.PR1 without CRISPR-Cas maintains five unique predicted prophages within its genome, similar to the number found in each *Nasonia*-associated isolate; and it also maintains the highest ANI to the *Nasonia* isolates at ∼92% as compared to the ∼84% of all other human isolates. The emphasis of these functions in human-associated *Providencia rettgeri* isolates, as opposed to insect-associated isolates, suggests human isolates are better poised to combat phage invasions.

To further tease apart the annotated functions specific in *Nasonia* and its close relative in *D. melanogaster*, we performed a functional enrichment analysis to statistically identify COGs only in the insect isolates **(Figure 3, Supplemental Table 4)**. The functional enrichment analysis computed in anvi’o uses a Generalized Linear Model with the logit linkage function to compute an enrichment score, p-value, and a False Detection Rate q-value between the different genome datasets [58, 59]. This analysis identified six COG functions unique to insect-associated genomes: L-rhamnose isomerase (COG4806), L-rhamnose mutarotase (COG3254), predicted metal-dependent hydrolase (COG1735), uncharacterized iron-regulated membrane protein (COG3182), putative heme iron utilization protein (COG3721), and Ca2+/H+ antiporter (COG0387) **(Supplemental Table 4)**. L-rhamnose is a common sugar component of plant and bacterial cell walls [69, 70], serves as a carbon source for some bacteria [71], and may confer protection to host immune proteins [72]. For example, adherent-invasive *E. coli* when grown on bile salts (a component of the gut) showed an upregulation of genes involved in sugar degradation, including the rhamnose pathway [73], and host-associated *L. monocytogenes* survival increases by glycosylating L-rhamnose and thus decreasing the cell wall permeability to antimicrobial peptides [72]. Additionally, as iron is a key metal for biological functions, this poses the question as to why these genes may be enriched in insect-associated environments as compared to those in human-associated settings. Iron-regulated membrane proteins and heme iron utilization proteins permit enteric bacteria to readily sense and respond to iron limiting environments and play a role in iron acquisition [74, 75]. Similarly, as mentioned above, the human-associated isolates encoded unique heme utilization proteins, emphasizing how these functions are important for specific host-associated environments. Insects differ from mammals in that they secrete ferritin, a protein that contains iron, into their hemolymph 1,000-fold higher than what is found in mammalian blood [76] and has altered expression in the presence of *Wolbachia* endosymbionts [77]. Perhaps the presence of these iron-regulating genes, particularly in insect-environments, helps to regulate iron homeostasis specific to what is available in insects. Lastly, the Ca2+/H+ antiporter may play a role in ion homeostasis to protect bacteria in altered pH environments and in maintaining internal pH homeostasis [78, 79]. It is important to note that although the unadjusted p-value for all genes in the enrichment analysis is significant (p=0.008), the adjusted q-value (q=0.897) that controls for false discovery rate does not reach significance due to the sample sizes for these genomes (n=7) **(Supplemental Table 4)**; however, this doesn’t change the presence/absence of these gene clusters across isolates.

#### Functional differences between Providencia rettgeri from Nasonia isolates versus humans and Drosophila

We were next interested in disentangling functions specific in the *Nasonia*-associated isolates. Therefore, we performed the same functional enrichment analysis to identify COGs that occur only in the *Providencia rettgeri* strains from *Nasonia* **(Figure 3, Supplemental Table 4)**. Only two COG categories occurred in the *Nasonia*-associated genomes: phospholipid N-methyltransferase (COG3963) and CYTH domain, found in class IV adenylate cyclase and various triphosphatases (COG2954) (p=0.03, q=1). Phospholipid N-methyltransferase synthesizes phosphatidylcholine (PC), a membrane-forming phospholipid present in only about 15% of bacteria [80, 81]. PC may help mediate symbiotic host-microbe interactions as PC is required for virulence in some bacteria and to establish beneficial symbioses with hosts [82]. Furthermore, adenylate cyclases are responsible for converting adenosine triphosphate (ATP) to cyclic adenosine monophosphate (cAMP) and play key regulatory roles such as mediating signal transduction in cells [83], of which the CYTH (CyaB, thiamine triphosphatase) domain of the type IV adenylate cyclases binds organic phosphate [84]. In *Pseudomonas aeruginosa*, when the adenylate cyclase gene CyaB was deleted, virulence was attenuated in a mouse model [85]. Collectively, these two COGs could play a role in mediating symbiotic relationships, beneficial or pathogenic, in the *Nasonia* host. Lastly, the Wasp.NvitL3.3 and Wasp.NgirL3.3 isolates did not share any mobile elements with any human or *Drosophila*-associated isolates. However, Fly.Dmel1 from *Drosophila* does share 62% nucleotide similarity with one phage spanning 46 genes from the previously published Wasp.Nvit03 *Nasonia* isolate (Phage 1; **Supplemental Figure 2**).

#### Functional differences within Providencia rettgeri Nasonia isolates

We further investigated the differences between *Providencia rettgeri* isolates within *Nasonia* species. Strains Wasp.NvitL3.3 and Wasp.NgirL3.3 isolated from the lab in Nashville, TN have a 99.99% ANI with each other and 99.3% ANI with Wasp.Nvit03 isolated from a lab located in Cambridge, MA **(Supplemental Figure 1)**; therefore, most of their genomic content is similar. No discernable difference could be identified between the two Nashville isolates; however, a small difference exists between these and the isolate from Cambridge derived from the same *N. vitripennis* line, AsymCx, less than six years apart **(Figure 3; Supplemental Table 2)**. *Providencia rettgeri* from Nashville encodes over three times as many genes relating to the COG category for [V] defense mechanisms (10.19% vs. 3.03% of total COG functions, **Supplemental Table 3)**. Although we found no evidence of CRISPR-Cas gene cassettes in *Nasonia*-associated genomes, a major difference is the presence of Type I restriction modification (RM) systems exclusively in the Nashville-associated *Providencia* genomes and absent from *N. vitripennis* strain Wasp.Nvit03 from Cambridge. Type I RM systems cleave away from the recognition site and have three components: a restriction enzyme (*hsdR*), a methyltransferase (*hsdM*), and a specificity subunit (*hsdS*) [86]. *Providencia rettgeri* strains Wasp.NvitL3.3 and Wasp.NgirL3.3 are predicted to encode two Type I RM systems, one with all three component genes and one missing the restriction enzyme. The Wasp.Nvit03 strain without the Type I RM system harbors six putative prophage regions within its genome, while both the newly sequenced Wasp.NvitL3.3 and Wasp.NgirL3.3 strains here contain three prophage regions each. The three phages found within Wasp.NvitL3.3 and Wasp.NgirL3.3 are 99.5-100% identical at the nucleotide level across the entire phage regions (**Supplemental Figure 2**), whereas the remaining three phage regions in Wasp.Nvit03 are novel or contain similarity to phage regions in *Providencia rettgeri* from *D. melanogaster*. The absence of both CRISPR-Cas and Type I R-M systems in the Wasp.Nvit03 strain could be directly correlated to the number of prophages within its genome, but a larger sample size of genomes may be necessary to make any firm predictions.

We also performed functional enrichment analysis to identify genes present only in *Providencia rettgeri* strains isolated from the lab in Nashville and absent in all other strains. We identified seven unique COGs: (i) CDP-glycerol glycerophosphotransferase (COG1887), (ii) uncharacterized conserved protein YBBC (COG3876), (iii) argonaute homolog implicated in RNA metabolism and viral defense (COG1431), (iv) predicted restriction endonuclease (COG3440), (v) DNA replication protein DnaD (COG3935), (vi) replicative superfamily II helicase (COG1204), and (vii) predicted ATPase archaeal AAA+ ATPase superfamily (COG1672) (p=0.03, q=1) **(Supplemental Table 4)**. Of particular interest is the Argonaute homolog and restriction endonuclease that function as viral and mobile element defense mechanisms [87, 88]. Recently, a bacterial Argonaute nuclease from *Clostridium butyricum* was shown to target multicopy genetic elements and suppress the propagation of plasmids and infection by phages via DNA interference [89]. In addition to the RM systems, this diverse suite of genes in *Nasonia* specifically from Nashville suggests enhanced protection against viral infection.

Conversely, strain Wasp.Nvit03 from Cambridge, MA encodes genes relating to type III secretion systems that are not present in strains from Nashville (COG4791, COG4790, COG4794) (p=0.03, q=1). Specifically, type III secretion systems serve to deliver effector proteins across bacterial and host membranes that can influence host cell biology [90]. For example, this machinery provides efficient protein transfer into eukaryotic cells that could inhibit phagocytosis or downregulate pro-inflammatory responses of the host [91]. Most notably, the biggest difference between the *Nasonia* isolates from Nashville vs. Cambridge is the unique phage machinery. Although two of the three prophages identified in Wasp.NvitL3.3 and Wasp.NgirL3.3 strains were also present in Wasp.Nvit03 based on amino acid identity, extensive rearrangements were evident **(Supplemental Figure 2)**. Notably, one additional phage in Wasp.Nvit03 has 73% nucleotide homology to a phage present in human isolate Human.RB151.

### *Proteus mirabilis* pangenomics

We employed the same methodology to compare the *Proteus mirabilis Nasonia*-associated genomes from this study (n=7) **(Table 2)** with high quality, publicly available genomes from close relatives in humans (n=8) **(Supplemental Table 1)**. We were interested in the diversity of *Proteus mirabilis* bacteria across related *Nasonia* species as they diverged one MYA, and their microbiomes exhibit strong signs of phylosymbiosis [24, 27, 30]. Importantly, *Proteus* bacteria exhibit low abundance in parental species, but F_2_ hybrids exhibit significant breakdown and lethality between the L3 and L4 larval stages; approximately 90% of the F_2_ hybrids die in conjunction with *Proteus* becoming the dominant bacteria that causes lethality [21]. Consequently, whether the *Proteus mirabilis* bacteria in the hybrids are the same or different from the parents remains a key question to understanding the nature of the bacteria-assisted hybrid breakdown. Our collection of *Proteus mirabilis* isolates encompasses two-three isolates each from *N. vitripennis, N. giraulti*, and F_2_ hybrid males to control for potential sex effects. Lastly, we also investigated how *Nasonia*-associated isolates compare to those isolated from human-associated environments. All *Proteus mirabilis* genomes are listed in Supplemental Table 1.

With all genomes combined (n=15), pangenomic analyses in anvi’o [58, 59] produced a total of 5,421 gene clusters. We grouped these gene clusters into three bins: (i) *Proteus* spp. Core (3,043 gene clusters), (ii) Human Specific (101 gene clusters), and (iii) *Nasonia* Specific (189 gene clusters) **(Figure 4a)**. Most gene clusters (56.1%) fall within the *Proteus* spp. Core bin where the majority of COG functions relate to [C] Energy production and conversion, [E] Amino acid transport and metabolism, and [J] Translation, ribosomal structure and biogenesis, the same three categories as reported for the *Providencia* species **(Supplemental Table 3, 5)**. Of the gene clusters not found in the core, 1.8% and 3.5% are shared just within human-associated and *Nasonia*-associated isolates, respectively, and the 38.5% remaining gene clusters are found within limited subsets of the genomes across isolates. Indeed, human-associated and *Nasonia*-associated *Proteus mirabilis* genomes share >= 98.6% average nucleotide identity (ANI) at the whole-genome level. Moreover, within *Nasonia*-associated isolates, the ANI increases to >= 99.9% identity **(Supplemental Figure 1)**. Collectively, this evidence demonstrates that *Nasonia Proteus mirabilis* are highly similar and slightly distinct from human *Proteus mirabilis*. Furthermore, using a set of 71 core bacterial proteins, we determined *Nasonia*-associated *Proteus mirabilis* are 99.9% identical and phylogenetically split from the human-associated strains **(Figure 2)**. Therefore, we next investigated what differences may exist between the genomes of (i) *Nasonia* species and hybrids and (ii) *Nasonia* and humans.

**Figure 4.**
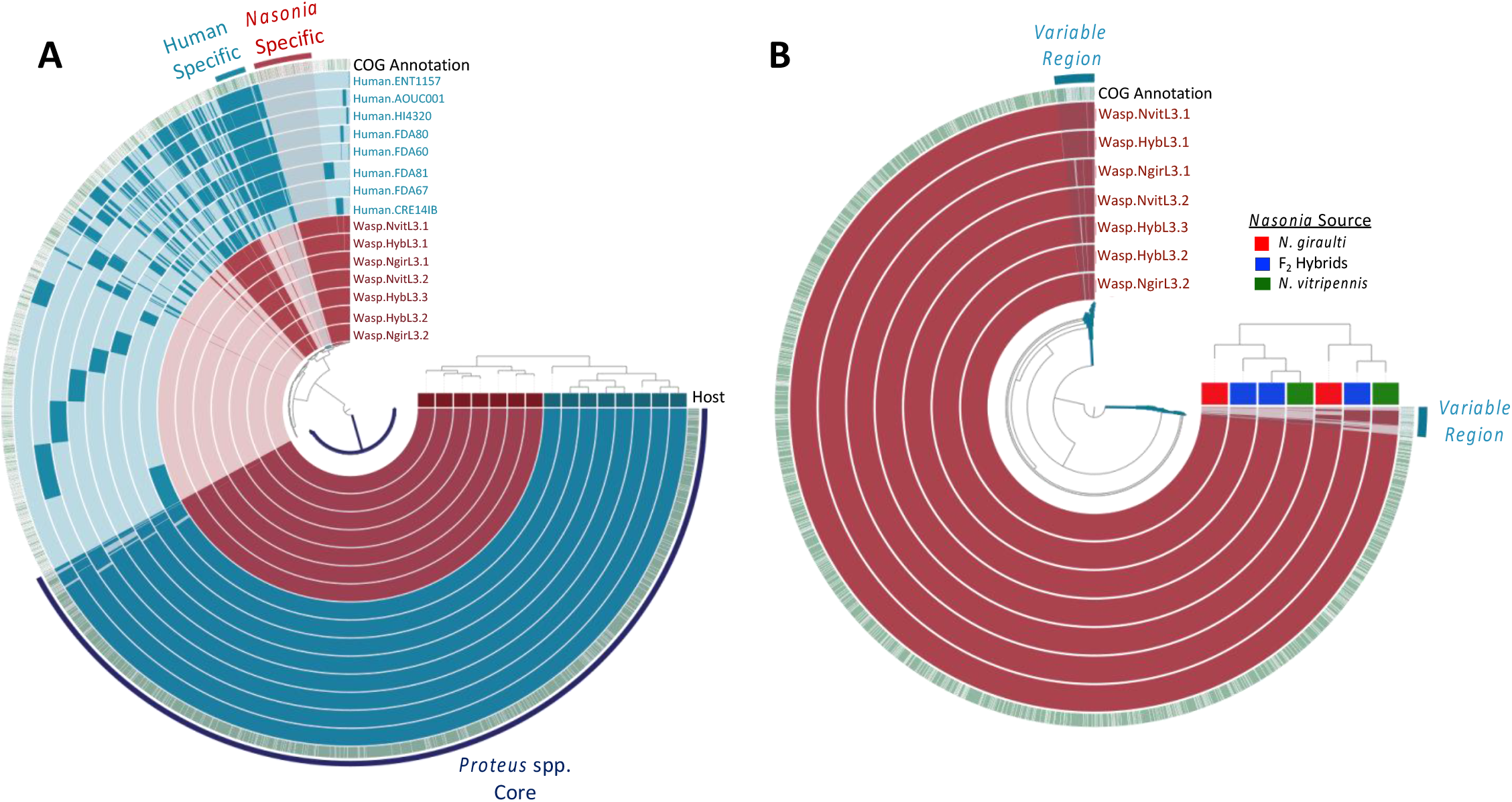
Pangenome of *Proteus mirabilis* human and *Nasonia* isolates. **A)** The fifteen inner layers correspond to all fifteen genomes, with *Nasonia* isolates shown in maroon and human isolates shown in blue. The top horizontal layer underneath the tree represent host source (*Nasonia* - maroon, Human - blue). **B)** The seven inner maroon layers correspond to the seven *Nasonia*-associated genomes sequenced as part of this study. The top horizontal layer underneath the tree represents host source (*N. giraulti* - red; F_2_ Hybrids - blue; *N. vitripennis* - green). For both **A** and **B**, the solid bars in each of the inner circles show the presence of gene-clusters (GCs) in a given genome while the outer green circle depicts known COGs (green) vs unknowns (white) assignments. The outermost layer of solid bars and text highlights groups of GCs that correspond to the genome core or to group-specific GCs. Genomes are ordered based on their gene-cluster distribution across genomes, which is shown at the top right corner (tree). Central dendrograms depict protein cluster hierarchy when displayed as protein cluster frequency.

#### Functional differences between Proteus mirabilis Nasonia isolates

We investigated the strain-level diversity within our *Nasonia*-associated *Proteus mirabilis* isolates to distinguish whether or not the *Proteus* strains in hybrid offspring are the same as those in parental *Nasonia*. Pangenomic analyses produced a total of 3,699 gene clusters **(Figure 4b)**. Functional enrichment analyses found no significant difference between *Proteus mirabilis* isolated from either *N. vitripennis, N. giraulti*, or F_2_ hybrids. Therefore, the hybrid *Proteus mirabilis* genomes are functionally identical to the parental *Proteus mirabilis* isolates. We identified two regions where there was variability in the distribution of gene clusters that totaled <0.05% of total gene clusters **(Figure 4b)** and grouped them into a bin labeled “Variable Regions” for further investigation. Inspection of this bin identified 92 gene clusters that occurred in subsets of the *Nasonia*-associated *Proteus mirabilis* genomes, of which 68 (74%) are lacking known functional annotations. No consistent trend emerged regarding gene clusters found only within specific *Nasonia*-associated genomes (e.g., gene clusters were not consistently unique in only *N. vitripennis* and hybrid strains or *N. giraulti* and hybrid strains). Hierarchical clustering based on gene cluster frequencies places the F_2_ hybrid strains as more similar to paternal *N. vitripennis* strains based on their gene-cluster distribution across genomes, but *N. giraulti* strains are still closely associated within these relationships **(Figure 4b)**, signifying that these small number of differences are not enough to designate the specific parental origin of the F_2_ hybrid strains.

For three phages (Phage 1-3), there is 100% nucleotide similarity and identical gene synteny across all seven *Proteus mirabilis* isolates in *Nasonia*, however key differences including gene deletions and truncations are apparent in proteins within the five other phages also present **(Supplemental Figure 3)**. These five phages have a nucleotide similarity ranging from 93.6-99.9% across the genomes, but the most significant dissimilarities are found within the phage tail regions, the part of the phage that directly interacts with its bacterial host and often determines host specificity. For example, in Phages 4 and 5 from Wasp.NgirL3.1, an early stop codon results in a predicted truncation in the tail spike protein, whereas across all other isolates containing these phages, the tail spike protein is intact. Relative to other isolates, Phage 6 found in Wasp.NvitL3.2 and Wasp.HybL3.2 both maintained the same two nucleotide deletions (4-mer and 8-mer) as well as mutations surrounding these deletions in the encoded host specificity protein J gene. Altogether, comparative genomics of the phage coding regions demonstrate that these isolates are extremely similar, and some of their phages exhibit distinct differences in the tail regions known to evolve fast, namely those that determine host specificity [92].

#### Functional differences between Proteus mirabilis Nasonia and human isolates

Next, comparisons between the *Nasonia* and Human bins **(Figure 4a)** revealed that the *Nasonia*-associated *Proteus mirabilis* are particularly enriched for gene clusters with functions for [M] cell wall/membrane/envelope biogenesis (14.58% vs 1.59%) **(Supplemental Table 3)**. This includes gene functions for glycosyltransferases involved in LPS biosynthesis (COG3306), peptidoglycan/LPS O-acetylase OafA/YrhL containing acyltransferase and SGNH-hydrolase domains (COG1835), and UDP-galactopyranose mutase (COG0562). LPS is a major component of Gram-negative bacterial cells walls that can be modified by glycosotransferases and acyltransferases, and variations in the structure of LPS can provide selective advantages in different environments such as inhibiting bacteriophage binding, temperature tolerance, and antimicrobial resistance [93–96]. Functional enrichment analyses in anvi’o identified the UDP-galactopyranose mutase COG function as only present in *Nasonia*-associated genomes (q=0.0173) and is responsible for the biosynthesis of galactofuranose, a sugar structure found in bacterial cell walls that is absent in mammals [97, 98] **(Supplemental Table 6)**. UDP-galactopyranose was shown to be essential for growth of *Mycobacterium tuberculosis* [99], although it remains unclear what function it may serve specifically in *Nasonia*-associated environments. Additionally, all *Nasonia*-associated genomes and a single human-associated genome (HI4320) encode muramidase (urinary lysozyme) (COG4678; q=0.0452), although a similar homologous COG function (COG3772) exists in all human-associated genomes and in three *Nasonia*-associated genomes as well. Murimidase activity has been directly connected to the ability for *Proteus* to differentiate from vegetative to swarmer cells [100], a phenotype that is visible from bacteria isolated here **(Figure 1b)**.

We also investigated putative adaptive and innate bacterial defense systems within the isolates including CRISPR-Cas, restriction-modification (RM), and BacteRiophage Exclusion (BREX) systems. Although we found no evidence of CRISPR-Cas gene cassettes in any of our isolates, there were predicted Type I RM and BREX systems. The specificity subunit (hsdS) of the complete Type I RM had an identical amino acid sequence to hsdS proteins (>95% nucleotide similarity) found in two other *Proteus mirabilis* strains (PmSDC32 from red junglefowl; CNR20130297 from human, as of September 2020). The putative Type I BREX system [101, 102] component genes were intact without frameshifts or premature stop codons. Although the BREX mechanism of foreign DNA removal is not yet understood completely, unlike CRISPR-Cas and RM systems, the defense system does not involve restriction of foreign DNA [101, 102]. However, both RM and BREX systems utilize methylation patterns to determine self/non-self-recognition. The absence of CRISPR-Cas systems in *Proteus* has been documented previously [103] although the absence has not been linked to expanded presence of mobile elements. The presence of the newly-characterized BREX defense system in *Proteus mirabilis* could prevent phage integration, but the evolutionary history behind either the prophage integration events in *Proteus mirabilis* or the history of the BREX system are unknown. No publicly available *Proteus mirabilis* strains appear to have the same Type I RM (as determined by the conserved methyltransferase protein). *Proteus mirabilis* strains MPE5139 (CP053684.1) and Pm15C1 (KX268685) contain a similar BREX cassette (86% query coverage; 97.75% similarity), but strains placed phylogenetically close to the *Nasonia*-associated isolates such as PmSC1111 (CP034090) and BC11-12 (CP026571) do not. The acquisition of the BREX system after integration of the numerous phage genomes found within *Proteus mirabilis* could explain the apparent lack of phage inhibition, although we cannot say whether this is the case in the current study.

We next investigated the human-associated *Proteus mirabilis* bin and discovered it encodes more unique gene clusters for functions relating to [P] inorganic ion transport and metabolism (9.56% vs 0% in *Nasonia* Specific bin) **(Supplemental Table 3)**. These gene clusters encompass multiple COG functions relating to outer membrane receptors for iron (COG1629), ferrienterochelin, and colicins (COG4771) as well as coper chaperone proteins (COG2608) and oxidoreductases (NAD-binding) involved in siderophore biosynthesis (COG4693) **(Supplemental Table 5)**. In addition to exploring the bins, we also performed a functional enrichment analysis in anvi’o [58, 59] **(Figure 4a)**. This analysis identified that siderophore biosynthesis (COG4693), previously noted in the human-associated *Proteus*, occurs in 7 out of 8 of the human-associated genomes and is absent in all *Nasonia*-associated genomes (q=0.045) **(Supplemental Table 6)**. Interestingly, the previously noted outer membrane receptor for ferrienterochelin and colicins (COG4771) has upregulated expression in pathogenic *E. coli* strains as compared to commensals [104]. Although homologous functions (different proteins with the same functional assignment) for outer membrane receptor proteins (COG1629 and COG4771) are also found in *Nasonia*-associated *Proteus mirabilis* genomes, the presence of oxidoreductase involved in siderophore biosynthesis (COG4693) and increased occurrence and diversity of genes relating to inorganic ion transport and metabolism in human-associated environments suggests that the more pathogenic strains may maintain a variety of antigens that could result in persistent inflammation and infection. When infection ensues, metal homeostasis can change dramatically, and therefore pathogens must be able to compete for the limited metal availability [105]. Vertebrate hosts, including humans, can produce proteins such as the iron-sequestering lipocalin 2 to keep trace metals away from bacteria, which in turn creates a selective pressure for bacteria to evolve diverse iron-binding siderophores and a higher affinity towards binding trace metals for thriving in vertebrate environments [105–107]. For example, uropathogenic *E. coli* in humans use the siderophore enterobactin (of which ferrienterochelin is the iron complex) to resist lipocalin 2 [108]. Therefore, the value of these proteins in human-associated environments can assist pathogens in resisting nutritional immunity imposed by the host [105, 109, 110].

Human-associated *Proteus mirabilis* isolates also encode 5x as many transposases than *Nasonia*-associated *Proteus* isolates. Transposons permit the movement of DNA in and around the bacterial chromosome, which can help facilitate growth and adaptation of bacteria to their host environment [111, 112]. Using the same functional enrichment analysis approach described above, we identified just two gene functions that are found in all human-associated *Proteus mirabilis* isolates (n=8) and absent in all *Nasonia*-associated isolates (q=0.017): (i) the type IV secretory pathway VirB4 component, and (ii) a predicted nuclease of restriction endonuclease-like (RecB) superfamily, DUF1016 family **(Supplemental Table 6)**. Type IV secretion systems secrete virulence factors in which the VirB4 subunit acts as an ATPase and is essential for some bacteria to cause infection [113, 114]. Additionally, restriction endonucleases, such as RecB, cleave DNA at specific sites and act as defense mechanisms for bacteria against foreign DNA [115]. Altogether, these differences in genomic content in human-associated strains suggest they may be more predisposed to succeed in environments where selective pressure may drive strategies for scavenging host metal nutrients and genome changes related to competition with mobile genetic elements.

## CONCLUSION

Host-associated microbial communities often establish intimate and distinguishable relationships that assist host metabolism [116], development [117], behavior [118], immune maturation [119], among others. Moreover, cross-system trends occur such as phylosymbiosis wherein microbial community ecological relationships recapitulate the host phylogenetic relationships. [15]. The *Nasonia* parasitoid wasp genus is a model system with phylosymbiosis in both the bacterial and viral compositions, hybrid maladies associated with the microbiome, and bacterial cultivability that permits further investigation of the host-microbe interactions that mediate these processes. Two bacterial genera dominate the larval microbiome of these wasps in pure species and hybrids, *Proteus mirabilis* and *Providencia rettgeri*, and here we have genomically characterized them relative to each other and to isolates from invertebrate and vertebrate animal hosts. There are three important findings from this work. First, this study reports the genome sequencing of bacteria from hybrid hosts. Recent sequencing of hybrid deep-sea mussels showed that their symbionts were genetically indistinguishable from parental mussels [120], and this premise remains to be evaluated in other hybrid systems. Second, the bacterial species in insect-associated environments are not human contaminants as they differ from other host-associated environments and have adapted unique functions for survival that may more tightly regulate symbiotic relationships in insects. Third, the *Proteus mirabilis* genomic diversity is not unique between *Nasonia* species and hybrids, thus supporting a tenet of hologenomic speciation whereby the dominant bacterium in hybrids is a resident microbial taxon in parental species. Therefore, just as the same alleles of nuclear genes in parental species underpin lethality in hybrids, so do bacteria from parental species in the case of *Proteus mirabilis* and *Nasonia* hybrids. Whole genome sequencing of both host and microbial constituents of this association now permit a deeper understanding of the multi-omic interactions between resident members of the microbiome and the host, which in turn underpin phylosymbiosis and hybrid breakdown in these wasps.

## MATERIALS AND METHODS

### Nasonia rearing

*Nasonia* species remain interfertile, in the absence of incompatible *Wolbachia* infections, as a result of their recent evolutionary divergence [121], which allows us to take advantage of their haplodiploid sex determination to acquire F_2_ recombinant hybrid male offspring from virgin F_1_ mothers following F_0_ crosses. We used the *Wolbachia* uninfected lines *N. vitripennis* AsymCx, *N. giraulti* RV2x(u), and F_2_ hybrids from paternal AsymCx x maternal RV2x(u) crosses. *Nasonia* were reared under 25°C constant light on *Sarcophaga bullata* pupae. Hybrids were generated as previously described [21]. Briefly, we collected virgin females and males from each parental species during early pupal development. Upon eclosion, parental adults were crossed in single-pair matings. F_1_ females were collected as virgins in early pupal stages and serially hosted after eclosion every 48h on two *S. bullata* pupae to generate F_2_ haploid recombinant males (collected on third hosting). Parental strains were reared concurrently under identical conditions.

### Isolate cultivation and sequencing

*Proteus* and *Providencia* bacteria were isolated from L3 larval stages of male *Nasonia giraulti* RV2x(u), *N. vitripennis* AsymCx, and F_2_ hybrids (paternal AsymCx x maternal RV2x(u)), using Tryptic Soy Agar (TSA) plates (Difco) containing 1.5% Agar. For each wasp line, 10 larvae were collected, and surface sterilized with 70% ethanol for one minute, washed with 1X Phosphate-buffered saline (PBS), and resuspended in 20 uL of 1X PBS. The resuspended individuals were homogenized using sterile pestles and serial dilutions were plated on TSA plates. Colonies with distinct morphology were sub-cultured on fresh TSA plates to ensure isolation and then stored as glycerol stocks (50% glycerol) at −80° C for future characterization.

For cultivation and characterization, isolates were maintained on TSA solid agar plates or in TSA broth grown at 37°C, shaking at 130 rpms in liquid culture. For sequencing, multiple distinct isolates were selected from each sample group (*N. giraulti, N. vitripennis*, and F_2_ hybrids), and genomic DNA was extracted from 3 mL of overnight (TSB) broth culture using the ZR Duet DNA/RNA MiniPrep Plus kit (Zymo Research) following manufacturer’s protocol for the “suspended cells” option. Isolates were designated via the addition of a letter corresponding to the host animal’s species: G for *N. giraulti*, V for *N. vitripennis*, and H for hybrid. DNA was sent to the North Carolina State University’s Genomic Sciences Laboratory and sequenced using a single flow cell of the Illumina MiSeq platform to produce 2 x 250 bp paired-end reads.

### Biofilm Growth

Colonies from two isolates (NvitL3-1V and NvitL3-3V) from *N. vitripennis* were grown in overnight cultures of LB broth at 37°C and 130 RPM. The next day, cultures were diluted to an OD_600_ of 0.05 for each isolate, corresponding to a 5 x 10^5^ colony forming units per milliliter. A total of 1 milliliter was plated onto plastic cell culture 12-well plates in triplicate and incubated in a humid chamber overnight at 37°C. To prepare the co-culture inoculate, *Proteus* and *Providencia* were mixed 1:1 before adding a total of 1 milliliter to the wells. After 24 hours, the supernatant was removed and the biofilm stained and measured using established protocols [122]. Briefly, the culture was gently removed, and the plates were gently submerged into distilled water to removed unadhered cells. The plates were allowed to dry within a biosafety hood. The dried plates were then stained with 0.1% crystal violet for 10 minutes, washed three times in distilled water, and allowed to dry again under the hood. The stain retained was then resuspended in 30% acetic acid, and the OD_600_ was measured and reported. Values were graphed in GraphPad Prism 8, and statistical significance was determined using a Kruskal Wallis test with Dunn’s correction for multiple comparisons.

To quantitatively determine the co-culture composition of the *Proteus mirabilis* and *Providencia rettgeri* biofilm community we developed a qPCR assay using primers specific for unique single-copy genes in each respective genome for *P. mirabilis* (5’-GGTGAGATTTGTATTAATGG and 5’-ATCAGGAAGATGACGAG, annealing temperature 58°C) and *P. rettgeri* (5’-AACTCGGTCAGTTCCAAACG and 5’-CTGCATTGTTCGCTTCTCAC, annealing temperature 66°C). *Proteus* primers were designed for the ureR gene using a previously reported forward primer [123] and a new reverse primer. *Providencia* primers were designed based on a phage gene only found within the *Providencia* genomes. The biofilm experiment was repeated as described above except once the supernatant was removed, the adherent cells in the biofilm were recovered from the 1:1 co-culture in 1 milliliter of LB broth. The cells were then pelleted by spinning at 10,000g for 10 minutes and supernatant removed. DNA from the resulting cell pellet was extracted using the Gentra Puregene Tissue Kit (QIAGEN) according to manufacturer’s protocol and diluted to ∼10 nanograms/microliter each. Amplification was with BioRad iTaq Universal SYBR green supermix in a CFX96 real-time C1000 thermal cycler (BioRad, Hercules, CA) and each qPCR reaction was performed at the following thermal profile: 95°C for 3 minutes, followed by 40 cycles of 95°C for 15 seconds and the respective annealing temperature for each qPCR primer for 1 minute. Samples were calculated using a standard curve generated from dilutions of larger gene products amplified of the same genes for *P. mirabilis* (5’-GCGATTTTACACCGAGTTTC and 5’-ATCCCCATTCTGACATCCAA) and *P. rettgeri* (5’-CCGTTGTGTGTTTGGTATCG and 5’-GTAAGCTGCGTGGATTGGTT). Primer specificity was determined in-silico by BLASTing each primer sequence against each genome and by testing each primer pair on DNA from each bacterial species and observing no amplification either by PCR or qPCR. Isolates and materials are available upon request.

### Bacterial genome assemblies and annotations

All reads were trimmed using Trimmomatic v0.32 [124] and quality checked with FastQC [125]. Reads were further filtered using the Geneious v. 11.1.5 filter and trim workflow [126]. The reads of each isolate were assembled using SPAdes 3.13.0 [127], and the quality of each assembly was determined using QUAST [128], reads under 2kb were discarded. Genomes were annotated using PROKKA [129] and visualized in Geneious v11.0.3. Genome completion and contamination estimates were calculated using CheckM [48] and Average Nucleotide Identity (ANI) using FastANI [55] online in Kbase (www.kbase.us) [130]. Prophages were identified using VirSorter [131], and plasmids were identified through PROKKA annotations and increased numbers of reads mapping through Bowtie2 [132]. All whole genome sequences were deposited in GenBank under BioProject PRJNA660265. BioSample accessions and further metadata are provided in Supplemental Table 1.

### Genomic comparisons

For phylogenetic placement of *Proteus* and *Providencia* bacteria isolated in this study, we aligned nucleotide sequences of the gyrase B subunit (*gyrB)* and RNA polymerase B subunit (*rpoB*) separately within Geneious 11.1.5 [126] using the multiple align tool. Using these alignments, we constructed separate unrooted maximum likelihood (ML) phylogenies with 1,000 replicates for bootstrapping, using the GTR GAMMA model within RAxML for *gyrB* from both *P. rettgeri* and *P. mirabilis*. Evolutionary models were determined with jModelTest [133]. Trees were visualized in FigTree v1.4.4 (http://tree.bio.ed.ac.uk/software/figtree/).

Concatenated amino acid sequences for phylogenomic analyses were computed in anvi’o [58] using ‘anvi-get-sequences-for-hmm-hits’ with the flags --return-best-hit, --get-aa-sequences, and –concatenate with the --hmm-source Bacteria_71: [type: singlecopy] ADK, AICARFT_IMPCHas, ATP-synt, ATP-synt_A, Adenylsucc_synt, Chorismate_synt, EF_TS, Exonuc_VII_L, GrpE, Ham1p_like, IPPT, OSCP, PGK, Pept_tRNA_hydro, RBFA, RNA_pol_L, RNA_pol_Rpb6, RRF, RecO_C, Ribonuclease_P, Ribosom_S12_S23, Ribosomal_L1, Ribosomal_L13, Ribosomal_L14, Ribosomal_L16, Ribosomal_L17, Ribosomal_L18p, Ribosomal_L19, Ribosomal_L2, Ribosomal_L20, Ribosomal_L21p, Ribosomal_L22, Ribosomal_L23, Ribosomal_L27, Ribosomal_L27A, Ribosomal_L28, Ribosomal_L29, Ribosomal_L3, Ribosomal_L32p, Ribosomal_L35p, Ribosomal_L4, Ribosomal_L5, Ribosomal_L6, Ribosomal_L9_C, Ribosomal_S10, Ribosomal_S11, Ribosomal_S13, Ribosomal_S15, Ribosomal_S16, Ribosomal_S17, Ribosomal_S19, Ribosomal_S2, Ribosomal_S20p, Ribosomal_S3_C, Ribosomal_S6, Ribosomal_S7, Ribosomal_S8, Ribosomal_S9, RsfS, RuvX, SecE, SecG, SecY, SmpB, TsaE, UPF0054, YajC, eIF-1a, ribosomal_L24, tRNA-synt_1d, tRNA_m1G_MT). Note: *Providencia rettgeri* strain Dmel1 was missing the gene for AICARFT_IMPCHas so this was manually removed from the *Providencia* alignment in Geneious resulting in a total of 70 genes. Concatenated alignments were imported into Geneious (https://www.geneious.com) [126] v11.0.3 and protein trees built with PhyML using an LG substitution model and 100 bootstraps. Evolutionary models were determined with Modeltest-ng for amino acids [133].

We used anvi’o [58] v6.2 following the pangenomics workflow [59] to analyze pangenomes of our *Proteus* and *Providencia* isolates with publicly available reference genomes from the National Center for Biotechnology Information (NCBI) (**Supplemental Table 1**), respectively. If genomes contained plasmids, plasmids were included within the genome file. Briefly, we used the program ‘anvi-script-FASTA-to-contigs-db’ to convert genome fasta nucleotide files into contigs databases for each genome which uses Prodigal [134] v2.6.2 for gene calling. We then annotated each gene using ‘anvi-run-ncbi-contigs’. A genomes storage file was created to collect each genome database using ‘anvi-gen-genomes-storage’ and the pangenome computed using ‘anvi-pan-genome’ with flags --mcl-inflation 10 and --use-ncbi-blast which uses the MCL algorithm [60, 61] to identify clusters in amino acid sequence similarity search results and blastp [135] for the amino acid sequence similarity search. Genomes were classified by host (human vs insect) using ‘anvi-import-misc-data’ and functional enrichment analyses were performed using ‘anvi-get-enriched-functions-per-pan-group’ with --annotation-source COG_FUNCTION. We defined “core” genes of each species pangenome as gene clusters that were present in every genome and accessory genes as those present in only a subset of genomes (e.g. *Nasonia* specific gene clusters). Figures were visualized in anvi’o interactive interface, Inkscape version 1.0 (available from https://inkscape.org/), and finalized in Microsoft Office PowerPoint (v16.37).

## Acknowledgments

This work was supported by the Vanderbilt Microbiome Initiative to S.R.B., National Institutes of Health Ruth Kirschstein Postdoctoral Fellowship to B.A.L., Deutsche Forschungsgemeinschaft (DFG) postdoctoral fellowship (MI 2242/1-1) to A. Mikaelyan, and Vanderbilt University SyBBURE Searle Undergraduate Research Program to A.K.M. Any opinion, conclusions or recommendations expressed in this material are those of the authors(s) and do not necessarily reflect the views of the National Institutes of Health, the National Science Foundation, or Vanderbilt University.

## Author Contributions

A.M. and S.R.B. conceptualized the study and designed experiments. A.M. performed microbial isolation and genome sequencing. K.L.C., B.A.L., E.A.H., and A.K.M. carried out experiments and performed data analysis. K.L.C. and B.A.L. analyzed the genomes and performed pangenomic analyses. K.L.C. and S.R.B. wrote the manuscript with input from B.A.L. S.R.B. supervised the project.

## FIGURES

**Supplemental Figure 1.**
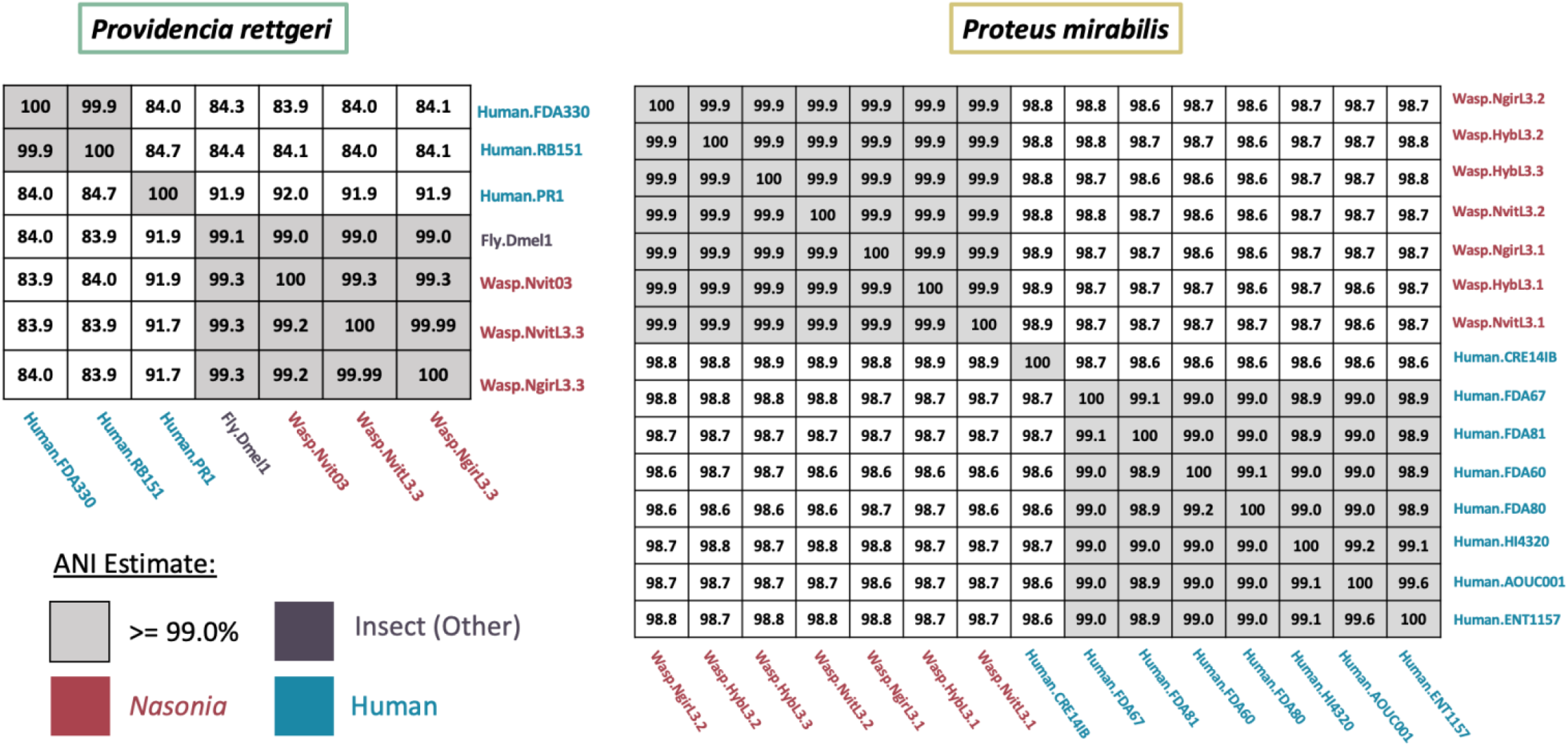
Whole genome average nucleotide identity comparisons between *Nasonia* isolates from this study and the publicly available genomes used for comparative genomic analyses.

**Supplemental Figure 2.**
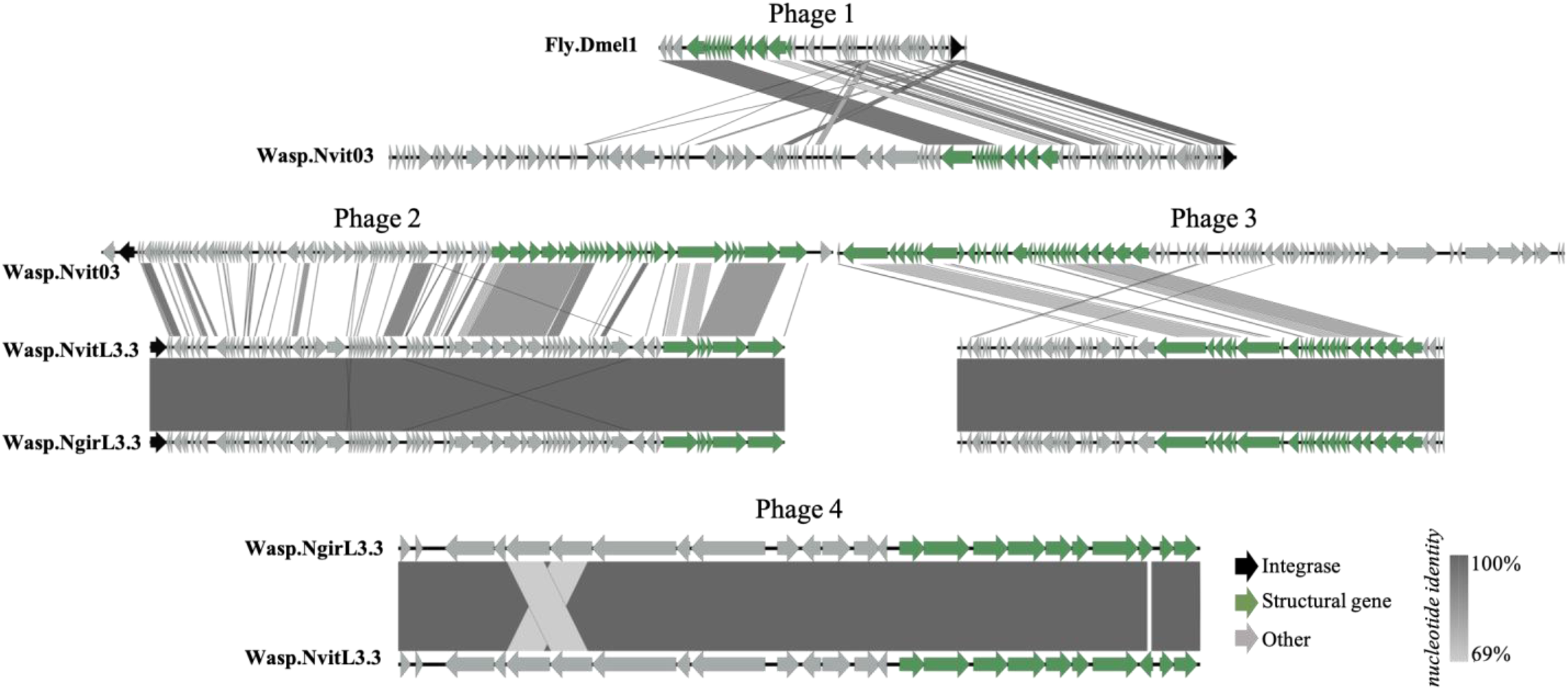
Phages of *Providencia rettgeri* insect isolates. Four total phages were identified among the insect isolates. Homologous phages were determined through whole genome alignments of putative prophages identified by VirSorter. Percent nucleotide identity as determined by blastn is indicated by shaded bars between each predicted coding gene.

**Supplemental Figure 3.**
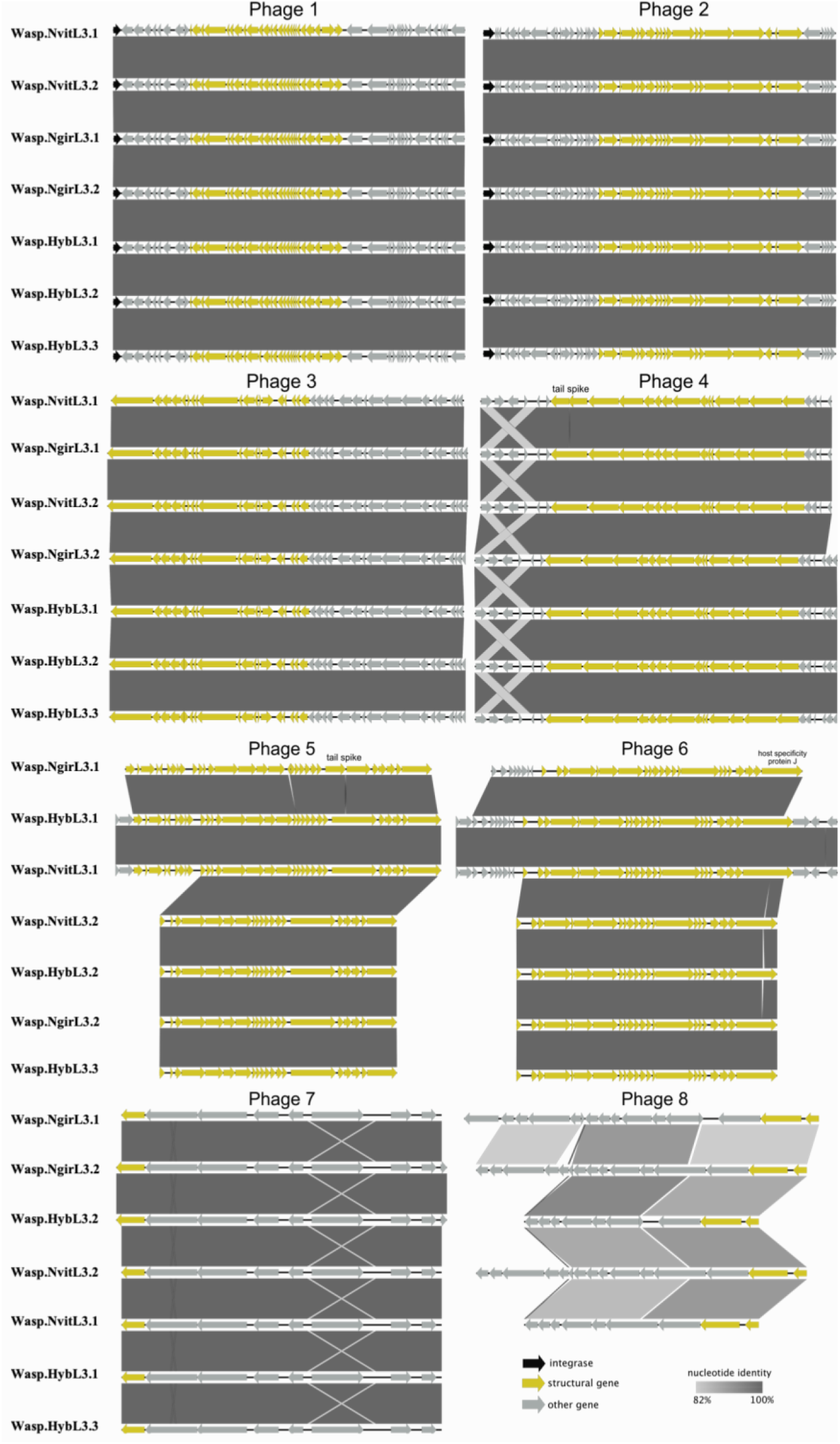
Phages of *Proteus mirabilis Nasonia* isolates. Eight total phages were identified. Homologous phages were determined through whole genome alignments of putative prophages identified by VirSorter. Percent nucleotide identity as determined by blastn is indicated by shaded bars between each predicted coding gene. Disrupted tail proteins are labeled.

**Supplemental Table 1**. Whole genome sequences used for comparative genomics analyses.

**Supplemental Table 2**. Genome Bins associated with the *Providencia rettgeri* pangenomic analysis of the seven *Providencia rettgeri* genomes.

**Supplemental Table 3**. COG analyses for comparison between the different *Providencia* anvi’o bins and the different *Proteus* anvi’o bins.

**Supplemental Table 4**. *Providencia rettgeri* enrichment analysis.

**Supplemental Table 5**. Genome Bins associated with the *Proteus mirabilis* pangenomic analysis of the fifteen *Proteus mirabilis* genomes.

**Supplemental Table 6**. *Proteus mirabilis* enrichment analysis.

## Notes

### Competing Interest Statement

The authors have declared no competing interest.

